# Innervation of cranial muscles requires Mllt11/Af1q/Tcf7c function during trigeminal ganglion development

**DOI:** 10.1101/2024.02.16.580667

**Authors:** Nicholas W. Zinck, Danielle Stanton-Turcotte, Emily A. Witt, Marley Bloomers, Angelo Iulianella

**Author notes:** **Contributions:** NWZ sectioned, immunostained, analyzed the data, generated figures, and wrote the first draft. MB and AI assisted with microscopy. DST and EAW helped with tissue preparation, mouse embryo generation, expression analysis, genotyping, and figures. AI supervised the project, obtained funding, conducted whole mount immunostaining and microscopy, edited figures and manuscript. **Funding:** Canadian Institutes of Health Research (CIHR PJT-388914), National Science and Engineering Research Council of Canada (RGPIN 03925-20).

## Abstract

The development of cranial nerves, including the trigeminal nerve, and the formation of neuromuscular junctions (NMJs) are crucial processes for craniofacial motor function. Mllt11/Af1q/Tcf7c (henceforth Mllt11), a novel type of cytoskeletal-interacting protein, has been implicated in neuronal migration and neuritogenesis during central nervous system development. However, its role in peripheral nerve development and NMJ formation remains poorly understood. This study investigates the function of Mllt11 during trigeminal ganglion development and its impact on motor innervation of the masseter muscle. We report Mllt11 expression in the developing trigeminal ganglia, suggesting a potential role in cranial nerve development. Using a conditional knockout mouse model to delete *Mllt11* in Wnt1-expressing neural crest cells, we assessed trigeminal ganglion development and innervation of the masseter muscle in the jaw. Surprisingly, we find that *Mllt11* loss does not affect the initial formation of the trigeminal ganglion but disrupts its cellular composition, with a reduction in the ratio of neural crest-derived Sox10^+^ cells relative to placode-derived Isl1/2^+^ cells. Furthermore, our study demonstrates that conditional *Mllt11* knockout leads to reduction of neurofilament density and NMJs within the masseter muscle, indicating altered trigeminal motor innervation. Our findings show that Mllt11 regulates the cellular composition of the trigeminal ganglion and is essential for proper trigeminal motor innervation in the masseter muscle.

## INTRODUCTION

The peripheral nervous system is composed of the cranial and spinal nerves, which function in relaying sensory and motor signals between the central nervous system and somatic tissues. The peripheral nerves are derived from the neural crest cells (NCCs) and cells from specialized regional thickenings of the cranial ectoderm called placodes (D’Amico-Martel & Noden 1983, Hamburger 1961, Koontz et al 2023, Trainor 2014). During peripheral nerve neurogenesis, both neural crest cells and ectodermal placode cells delaminate from their respective epithelia, interact and ultimately lead to the formation of the peripheral nerves (Hamburger 1961, Kurosaka et al 2015). While peripheral nerve development has been a subject of study for many decades, the process is not fully understood.

The cranial nerves (CNI-CNXII) are a subset of peripheral nerves originating from the supraspinal elements of the central nervous system. In mice, the cranial nerves emerge at approximately embryonic day (E) 9.5, and axonal projection is apparent by E10.5 (Kurosaka et al 2015, Sudiwala & Knox 2019). Each of the twelve cranial nerves provides either sensory, motor, or mixed sensory/motor function to the craniofacial and cervical regions, as well as various abdominothoracic viscera in the case of parasympathetic innervation via the vagus nerve (CNX) (Cordes 2001). The variety and importance of functions regulated by the cranial nerves is highlighted by the various disorders associated with cranial nerve dysinnervation, collectively known as *cranial nerve dysinnervation disorders*. In the literature, special attention has been paid to disorders characterized by dysinnervation of the oculomotor (CNIII), trochlear (CNIV), trigeminal (CNV) and, abducens (CNVI) nerves due to their importance in ocular and facial motor function (Kaeser & Brodsky 2013, Traboulsi 2004). A further understanding of cranial nerve development is crucial to understand the pathophysiology of cranial nerve dysinnervation disorders.

Cytoskeletal formation and remodeling are critical processes in both neurogenesis and neuritogenesis (i.e., the formation of axons and dendrites), enabling cell migration, differentiation, and neurite growth (da Silva & Dotti 2002). We previously characterized Myeloid/lymphoid or mixed-lineage leukemia; translocated to chromosome 11 or ALL1 fused from chromosome 1q or T cell factor 7 cofactor (Mllt11/AF1q/Tcf7c; thereafter referred to as Mllt11) as a vertebrate-specific cytoskeletal-associating protein with restricted expression in the developing central nervous system by E9.5 and in the peripheral ganglia from E10.5 onwards (Yamada et al 2014). More recently we showed that Mllt11 is required for neuritogenesis of cortical neurons both *in vitro* and *in vivo*, with *Mllt11* mouse null mutants exhibiting markedly reduced cortical neurite length and complexity, which affected the formation of cortical white matter tracts and commissural projections (Stanton-Turcotte et al 2022). In the same study we also showed that *Mllt11* loss-of-function resulted in reduced migration of neurons towards the cortical plate. Mllt11 has also been implicated in cell migration and morphogenesis during retinogenesis. Specifically, knockout of *Mllt11* led to failed migration of retinal ganglion cells and amacrine cells during retinal lamination, and these cells were unable to elongate and assume a bipolar morphology consistent with migrating neuroblasts and instead adopted a more globular morphology associated with reduced migratory ability (Blommers et al 2023). While we previously showed that Mllt11 regulates neuronal migration and neuritogenesis in the central nervous system, its role in the peripheral nervous system has yet to be explored. Using a *Wnt1*^*Cre2/+*^ approach to inactivate a floxed *Mllt11* allele in NCC derivatives, we now report a requirement for Mllt11 in the formation of the trigeminal nerve, the largest cranial nerve, and innervation of the masseter muscle. This is consistent with a role for Mllt11 in neuritogenesis of all neurons, irrespective of their embryonic origins.

## METHODS

### Animals

All experiments were done according to approved protocols from the IACUC at Dalhousie University. *Mllt11*^*flox/flox*^*;Rosa26r*^*tdTomato/tdTomato*^ (Ai9; Jackson Laboratory strain#00709) reporter mice were generated and genotyped as previously described (Stanton-Turcotte et al 2022). Neural crest cell specific ablation of *Mllt11* was accomplished by mating *Wnt1*^*Cre2/*+^ driver mice (B6.Cg-E2f1Tg(Wnt1-cre)2Sor/J; Jackson laboratory strain#022501) mated to *Mllt11*^*flox/+*^*;Rosa26r*^*tdTomato/tdTomato*^ to generate NCC-specific conditional knockout used in this study. To generate neural crest cell specific ablation of *Mllt11*, the *Wnt1*^*Cre2/*+^ allele was derived maternally, as previously reported (Dinsmore et al 2022). Confirmation of trigeminal ganglion Mllt11 expression was done using the targeted pre-floxed *Mllt11* allele, which houses a β-galactosidase cassette (*Mllt11*^*tm1a(KOMP)Mbp*^) (Stanton-Turcotte et al 2022).

### Histology

Embryos were dissected from maternal decidua and either whole embryos (E12.5) or heads (E18.5) were fixed in 4% paraformaldehyde in phosphate buffer for four hours depending, cryoprotected in sucrose graduation, and embedded in optimum cutting temperature (OCT) compound (Tissue-Tek, Torrance, CA). Embedded samples were stored at −80°C until sectioning. Samples were cryosectioned at 20 μm transversely and stored at −20°C until stained. ß-gal staining was performed with the *Mllt11*^*tm1a(KOMP)Mbp*^ targeted allele (Stanton-Turcotte et al., 2022), using serial incubations with warmed ß-Galactosidase Stain Base Solution A and then Solution B, according to manufacturer’s instruction (Millipore, Burlington, MA), and staining with X-gal solution (Novagen, Madison, WI).

### Immunohistochemistry

Frozen sections were permeabilized by incubating in 0.1% Triton/Phosphate Buffered Saline with 0.1% Triton X-100 (PBT) and incubated in 3% normal donkey serum (NDS)/3% bovine serum albumin (BSA)/0.1% PBT for 1h at room temperature. The sections were then incubated with primary antibodies overnight at 4°C. Primary antibodies utilized include rabbit anti-Sox10 (1/100, Abcam), mouse anti-Isl1/2 (1/25, DSHB), mouse anti-Neurofilament 2H3 (1/200, DSHB), and rabbit anti-Cleaved Caspase-3 (1:500, Cell Signaling Technology). Alpha-bungarotoxin (BTX) conjugated with Alexa Fluor 647 (1:500, Invitrogen) was used in order to visualize nicotinic acetylcholine receptor densities. Species-specific Alexa Fluor 488-, 568-, and 647-conjugated secondary antibodies were used to detect primary antibodies (1:1500, Invitrogen). Following antibody staining, nuclei were counterstained with DAPI (4′,6-diamidino-2-phenylindole; 0.6mg/mL, Sigma). Following washes, slides were mounted (DakoCytomation fluorescent mounting medium) and stored at 4°C until imaging.

For whole mount immunostaining, embryos were dissected in Phosphate Buffered Saline (PBS), selected for tdTomato+ reporter (*Rosa26r*^*tdTomato/tdTomato*^) expression in *Wnt1*^*Cre2/+*^-fate mapped neural crest cell derivatives, fixed with 4% paraformaldehyde/PBS pH 7.4 for 4-5 hours at 4°C, rocking. Then embryos were rinsed in PBT, blocked in 3% Donkey Serum/1% Bovine Serum Albumin/PBT for 2-24 hours at 4°C. After genotyping embryos, controls (*Wnt*^*Cre2/+*^; *Mllt11*^*Flox/+*^; *Rosa26r*^*tdTomato/tdTomato*^) were pooled separately from *Mllt11* conditional mutants (*Wnt1*^*Cre2/+*^; *Mllt11*^*Flox/Flox*^; *Rosa26r*^*tdTomato/tdTomato*^) and incubated with mouse anti-β3 Tubulin (TuJ1, 1:1000, MilliporeSigma) antibodies overnight at 4°C, rocking. The following day, embryos were rinsed extensively in PBT, and incubated in donkey anti-mouse Alexa Fluor 488 secondary antibodies (1:1000, Invitrogen) overnight at 4°C, rocking. The following day embryos were rinsed in PBT, counterstained with DAPI, and cleared in glycerol/PBS buffer.

### Microscopy and Imaging

Microscopy was performed using a Zeiss AxioObserver fluorescence microscope equipped with an Apotome 2, 10x and 20x objectives, and Hamamatsu Orca Flash v4.0 digital camera. For whole mount immunostaining, microscopy was performed using a Zeiss V16 Zoomscope equipped with an Axiocam 506 mono digital camera. Images were processed using Zen software (Zeiss, Baden-Württemberg, Germany) and Affinity Photo 2 (Serif, Nottinghamshire, UK).

### Image sampling, quantification, and analysis

Cell counts were performed at E12.5 for DAPI, Sox10, Isl1/2^+^ cells, as well as *Wnt1*^*Cre*^; *tdTomato*^*+*^ fate mapped NCCs. Three regions of interest (ROI) 100 x 100 μm in size were randomly placed over images of immunostained sections of trigeminal ganglia and the number of cells within each ROI were counted using ImageJ (Fiji). For each individual animal, this was performed on three serial sections of the same ganglion. Cell counts were normalized over the number of DAPI^+^ cells in the same ROI (e.g., Wnt1^Cre2/+^;tdTomato^+^/DAPI^+^ cells) and averaged for each individual animal. These values are represented by data points in Figures 2, 3 and 4.

**Figure 1.**
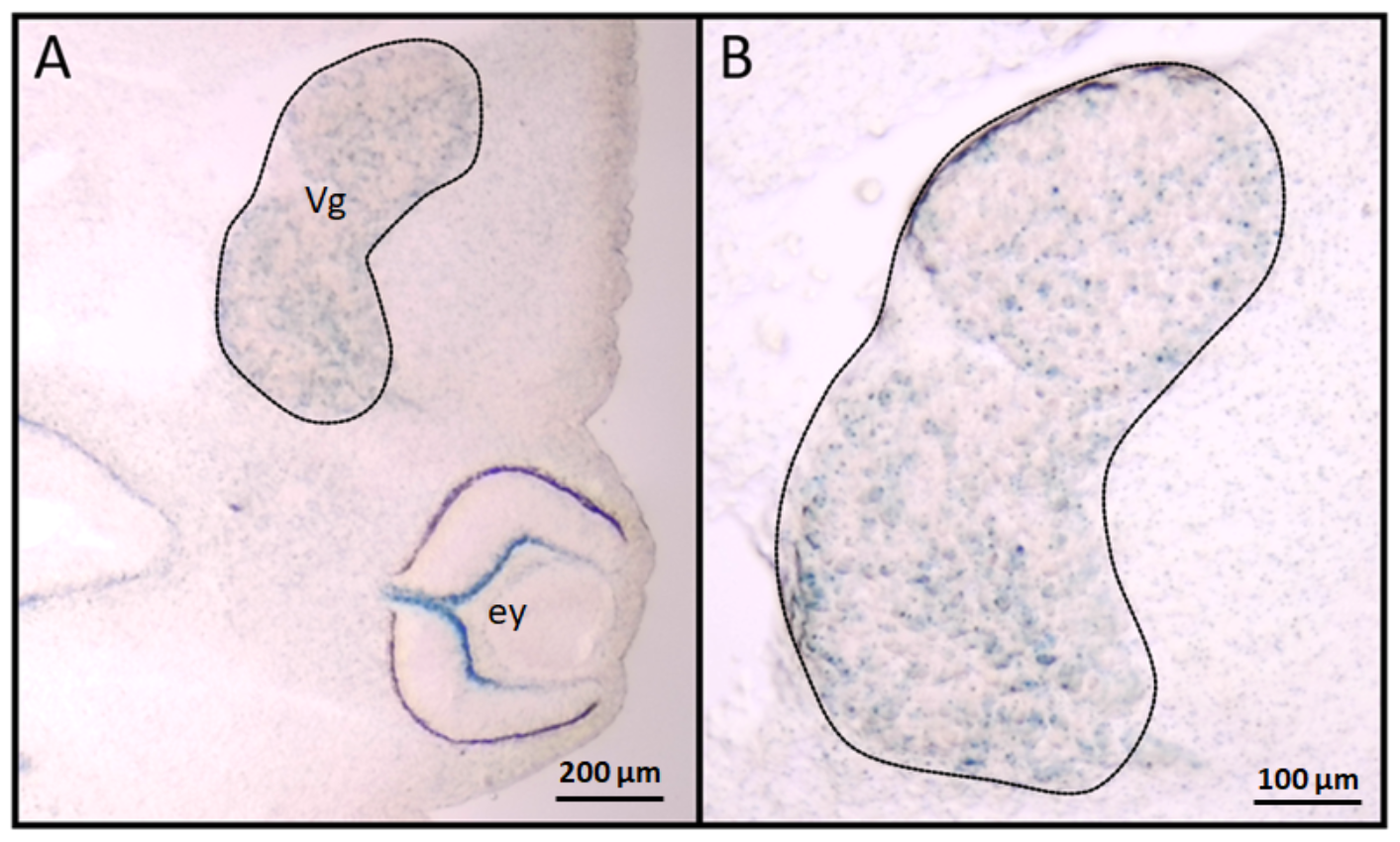
*Mllt11* is expressed in the trigeminal ganglion. *Mllt11* locus with a β-gal knock-in cassette reveals expression in the trigeminal ganglion at E12.5. Arrow indicates trigeminal ganglion (A). Blue represents X-gal precipitate associated with Mllt11 expression.

**Figure 2.**
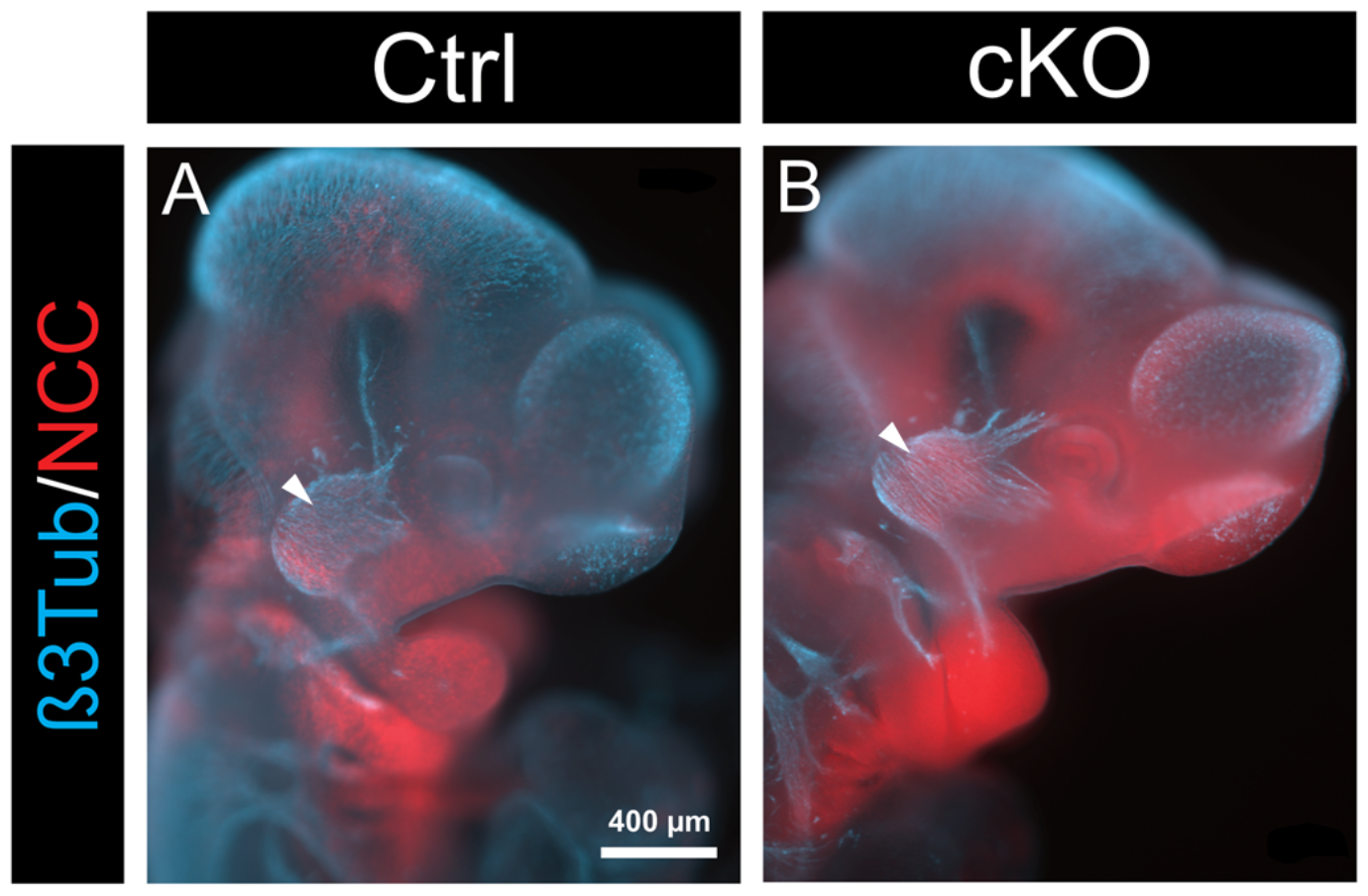
Normal trigeminal (Vg) nerve morphology in *Mllt11* mutants. Whole-mount immunohistochemistry for ß3Tub at E10.5 illustrating normal Vg cranial nerve morphology in *Mllt11* cKO mutants. Blue, ß3Tub; red, *Wnt1*^*Cre2/+*^*:Rosa26r*^*tdTomato/+*^ fate map; lateral view. Arrowhead indicates trigeminal ganglion.

**Figure 3.**
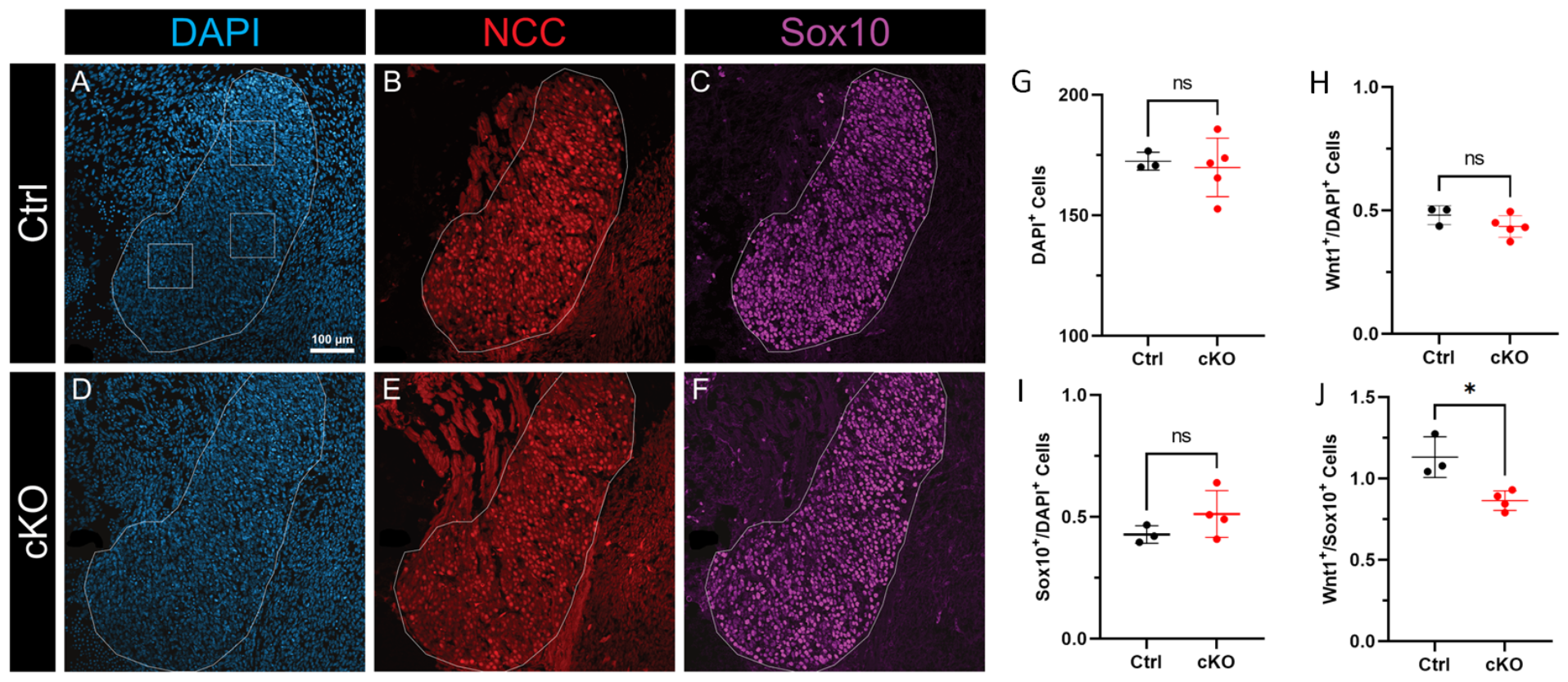
Reduced numbers of NCC-derived Sox10^+^ cells in *Mllt11* cKO mutants. Control and cKO E12.5 trigeminal ganglia positive for DAPI (A, D), NCCs (*Wnt1*^*Cre2/+*^; *Rosa26*^*tdTomato/+*^; B, E), and Sox10 (C, F) in transverse sections. Ganglia are outlined in white. Boxes represent ROIs for cell-count sampling. G, H, I, J, plot comparing the total cell density (G), proportion of NCC (H) and Sox10+ (I) cells, and the ratio of NCCs^+^ to Sox10^+^ cells (J) in the trigeminal ganglia between control (black) and cKO embryos (red). Error bars indicate standard deviation; ns, not statistically significant; * p < 0.05.

**Figure 4.**
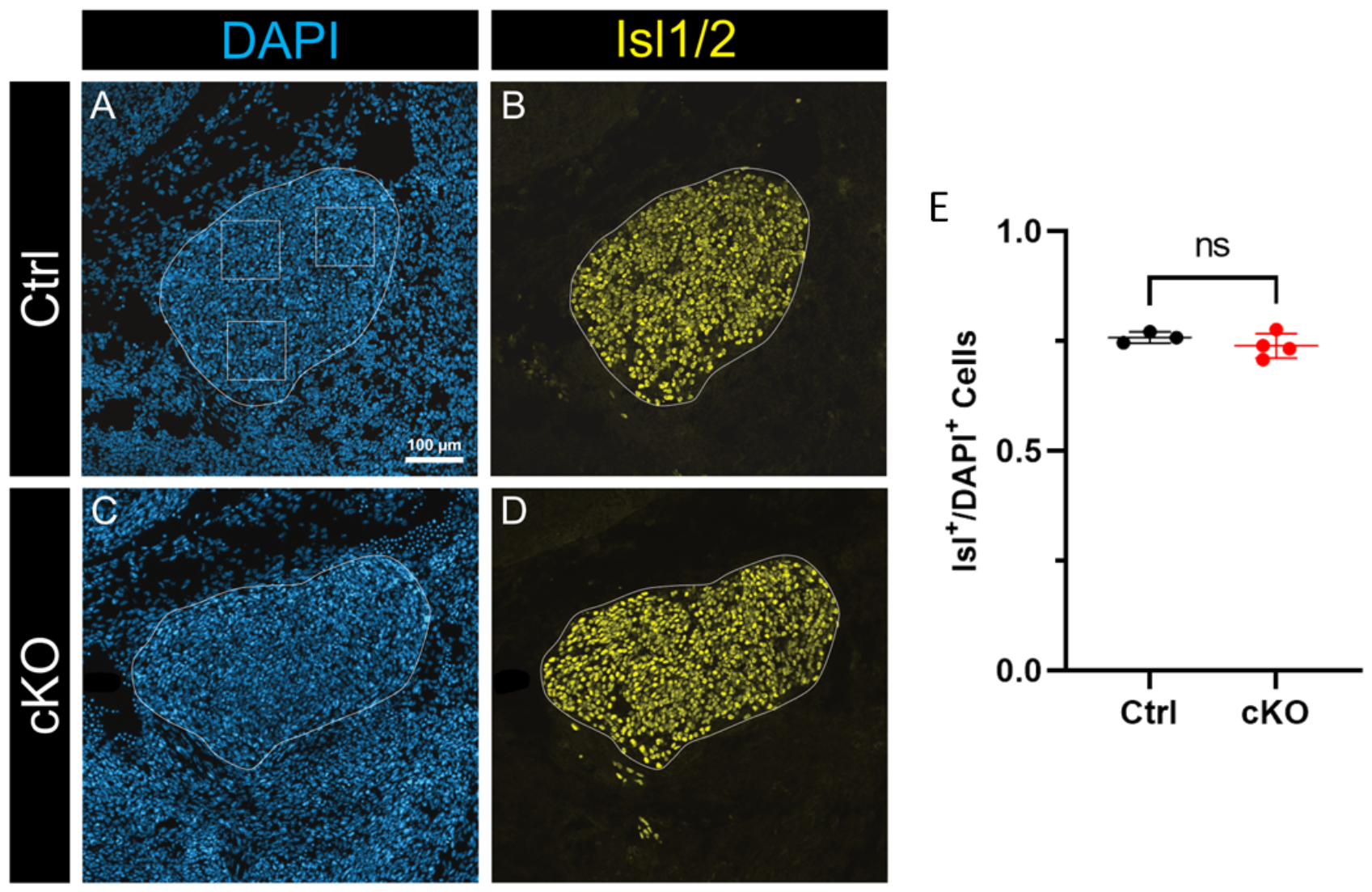
Placode-originating cells are unaffected in *Mllt11* cKO mutants. Control and cKO E12.5 Vg stained with DAPI (A, C), and anti-Isl1/2 (B, D). Ganglia are outlined in white. Boxes represent ROIs for cell-count sampling. E, plot comparing ratio of Isl1/2^+^ and DAPI^+^ cells between controls (black) and Mllt11 cKO (red). Error bars indicate standard deviation; ns, not statistically significant. Abbreviations: ROI, regions of interest; Vg, trigeminal ganglia.

Cell counts were conducted at E18.5 within a ROI comprised of the total area of masseter muscle due to the non-uniform nature of nerve distribution within the masseter. For each individual animal, this was performed on three serial sections of the same ganglion. Cell counts were normalized to the total area of the ROI and averaged for each individual animal. These values are represented by data points in Figures 6 and 7. An identical method was used for the analysis of CC3 signal in the E12.5 cell death assay (Figure 5).

**Figure 5.**
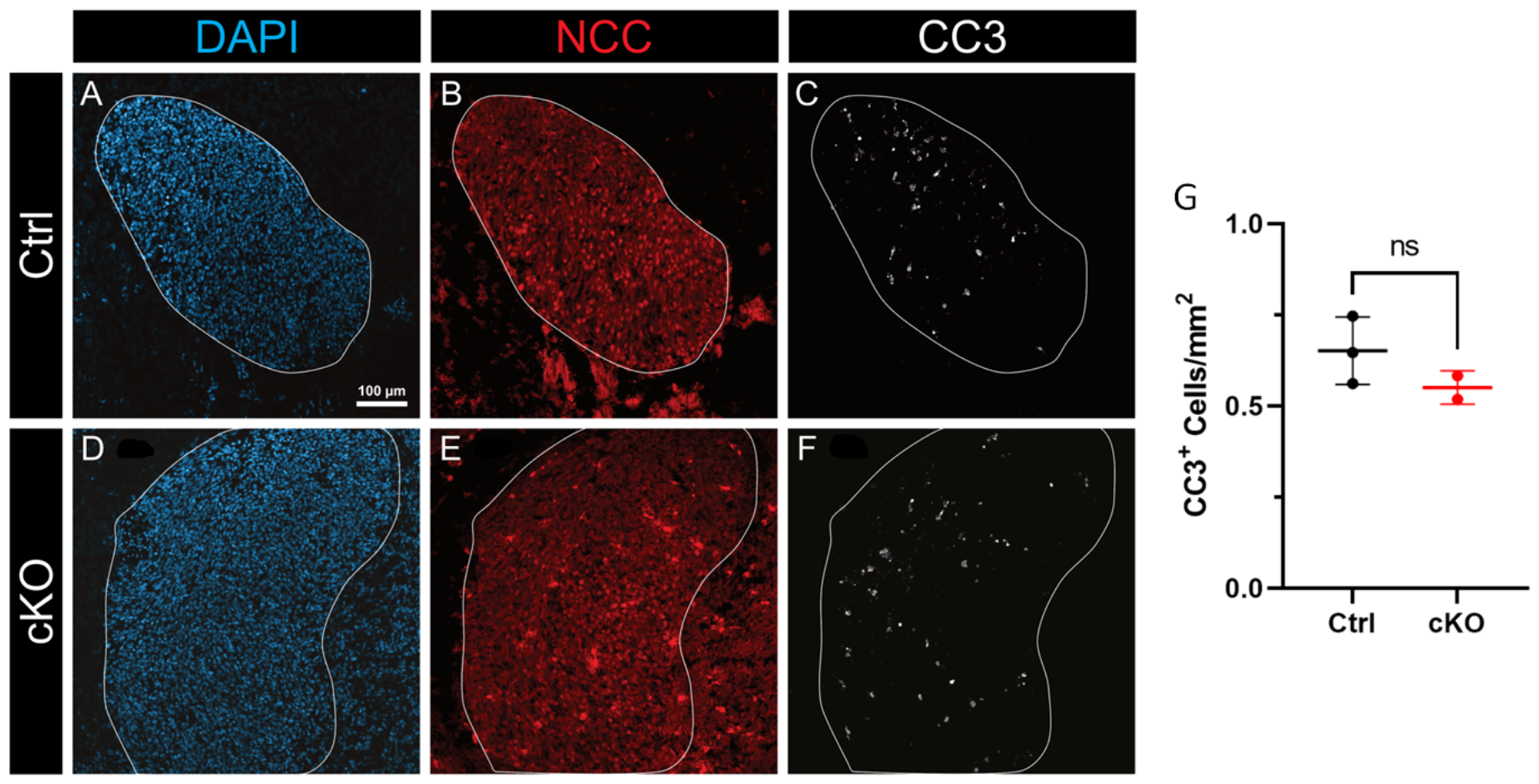
Trigeminal ganglion composition differences in *Mllt11* cKO mutant is not explained cell death. Control and cKO E12.5 trigeminal ganglia positive for DAPI (A, D), NCC (B, E), and Cleaved Caspase 3 (CC3; C, F) in transverse sections. Ganglia are outlined in white. G, plot comparing density of CC3+ cells in the E12.5 trigeminal ganglia between control (black) and cKO (red) mice. Error bars indicate standard deviation; ns, not statistically significant.

**Figure 6.**
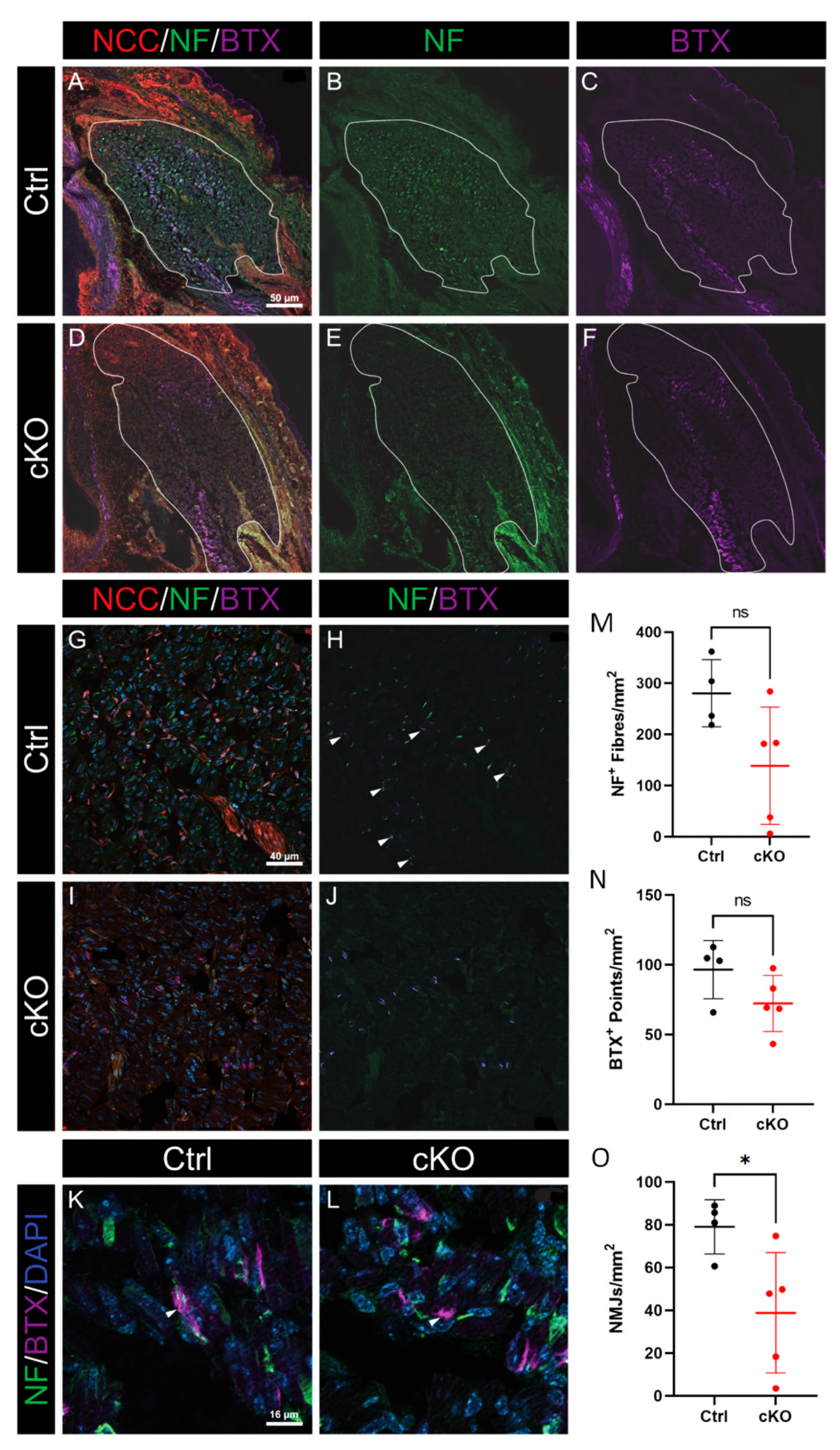
Masseter muscle innervation is disrupted in *Mllt11* cKO mice. Control and cKO E18.5 masseter muscles in transverse section displaying NCC (A, D, G, I), NF (A, B, D, E, G, I, K, L), BTX (nAchR; A, D, C, F, G, I, H, J, K, L), and DAPI (K, L) signals in transverse sections. Muscles are outlined in white. NMJs consisting of NF^+^ and BTX^+^ points in close contact, are indicated by white arrowheads (H, K, L; J lacks intramuscular NF signal). Representative images of intact (K) and disrupted NMJs (L). Plots comparing density of NF^+^ fibres (L), BTX^+^ points (M), and NMJs (N), all normalized over muscle area, between control (black) and cKO (red) mice. Error bars indicate standard deviation; ns, no statistically significant difference; *, p < 0.05.

## RESULTS

Previous studies have identified expression of Mllt11 in the cranial ganglia (Yamada et al 2014). In order to verify *Mllt11* activity in the developing trigeminal ganglia we utilized a β-gal knock-in allele for the *Mllt11* locus, as we previously described (Stanton-Turcotte et al 2022). Visualization of β-gal expression at E12.5 suggests that *Mllt11* locus is active within the trigeminal ganglion during development (Figure 1). We next sought to determine the role of *Mllt11* in the formation of the trigeminal ganglion and innervation of one of its main targets, the masseter muscle of the jaw. In order to investigate this, we generated *Wnt1*^*Cre2/+*^*;Mllt11*^*flox/flox*^*;Ai9 (Rosa26r*^*TdTomato/+*^) conditional knockout (cKO) embryos that had undergone targeted deletion of *Mllt11* driven by the *Wnt1*^*Cre2*^ transgenic allele to target NCC derivatives (Dinsmore et al 2022, Stanton-Turcotte et al 2022). We focused on the development of the trigeminal ganglion as it is the largest cranial ganglion that is significantly contributed by NCCs (Kurosaka et al 2015).

### *Mllt11* is not required for the formation of the trigeminal ganglia

We first confirmed *Wnt1*^*Cre2/+*^ transgene activity in the NCC neurogenic derivatives of the embryonic head, using *Cre*^*+*^ males to generate NCC-specific targeting, as recommended previously (Dinsmore et al 2022), by profiling Ai9 reporter activity. The resulting tdTomato^+^ signal identified NCC fate-mapped populations including the trigeminal nerve and its branches, which were visualized by ß-tubulin III (ß3Tub) whole mount immunostaining at E10.5. We observed no difference in overall trigeminal ganglion morphology (Figure 2), suggesting that the initial formation of the structure is unaffected by *Mllt11* loss. We next examined whether of *Mllt11* loss affected the cellular composition of the ganglia in the *Wnt1*^*Cre2/+*^-driven *Mllt11* cKO mice using lineage markers for cells arising from the neural crest (Sox10) vs. embryonic ectodermal placodes (Isl1/2) (Figures 3 and 4, respectively). At E12.5, there was no statistically significant difference in the mean number of cells within the ganglia (Figure 3A, D, G; Table 1; p=0.739, T-test). Additionally, there was no statistically significant difference in the proportion of *Wnt1*^*Cre2/+*^-fate labeled NCCs (Figure 3B, E, H; Table 1; p=0.184, T-test) or Sox10^+^ cells (Figure 3C, F, I; Table 1; p=0.215, T-test;) relative to DAPI^+^ cells populating the trigeminal ganglion. Interstingly, *Mllt11* cKO mice displayed a high degree of variation in total cell density (Figure 3G), as well as the proportion of tdTomato^+^ NCCs and Sox10^+^ cells, relative to controls (Figure 3H, I). Moreover, *Mllt11* cKOs displayed a reduction in the ratio of NCCs to Sox10^+^ cells within the trigeminal ganglion, compared to controls (Figure 3J; Table 1; p=0.013, T-test). We next probed for alterations in placodal populations of the trigeminal ganglion by compared the number of Isl1/2-expressing cells between *Mllt11* cKO mutants and control mice, but found no difference in the number of Isl1/2^+^ cells (Figure 4; Table 1; p=0.329, n=4, T-test). This suggested that *Wnt1*^*Cre2/+*^-driven *Mllt11* cKOs specifically affected NCC-derived populations of the ganglion, such as Sox10^+^ glial cells, and not placodal cells, which are instead derived from ectoderm.

**Table 1.**
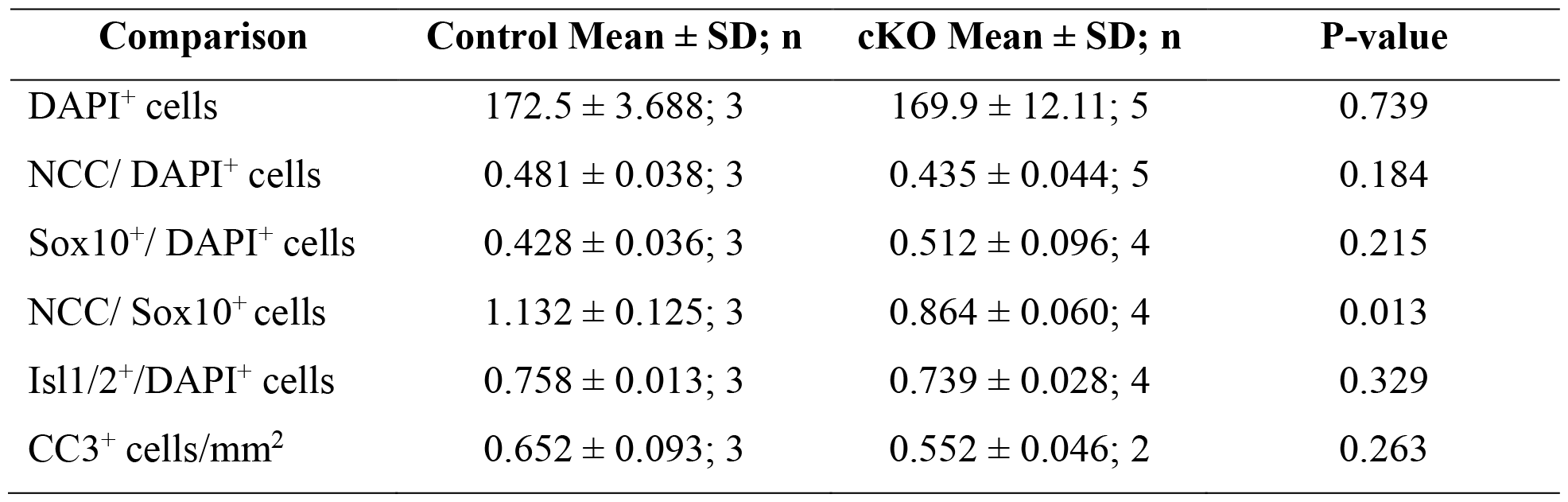
Comparisons of trigeminal ganglion cellular composition at E12.5. SD, standard deviation. NCCs were fate labeled with tdTomato^+^. P-value derived from T-test.

Given that *Mllt11* loss affected the NCC-derived cell numbers in the trigeminal ganglion, we evaluated if this was due to enhanced cell death by staining for Cleaved Caspase-3 (CC3) (Porter & Janicke 1999), but found no evidence of enhanced apoptosis in cKO ganglia (Figure 5; Table 1; p=0.263, T-test).

### *Mllt11* cKO in NCCs disrupts trigeminal motor innervation of the masseter

Our previous work demonstrated a role for Mllt11 in neuritogenesis and axonal targeting in the forebrain (Stanton-Turcotte et al 2022). As no major difference in trigeminal ganglion development was obvious at E12.5, we reasoned that NCC-specific ablation of *Mllt11* may lead to defects in the innervation of cranial muscles. We focused on trigeminal motor innervation and examined the presence of nerve fibres and neuromuscular junctions (NMJs) in the masseter muscle. We used neurofilament (NF) immunostaining along with neuromuscular endplate labeling with fluorescent bungarotoxin (BTX) staining, which identifies nicotinic acetylcholine receptors on muscle fibres. Interestingly, we observed a high degree of variation in NF density in the masseter of E18.5 *Mllt11* cKOs compared to controls (Figure 6B, E, H, J, M; Table 2; p=0.066, T-test), with the majority of individuals exhibiting reduced NF^+^ fiber staining in the masseter relative to controls (Figure 6M). In contrast to the disruption in NF bundle density, we did not observe a difference in the density of BTX^+^ staining of NMJs between control and cKO mice (Figure 6C, F, H, J, N; Table 2; p=0.120, T-test). Under higher magnification, we confirmed a striking reduction in the number of NMJs (NF^+^/BTX^+^) in E18.5 *Mllt11* cKO mouse masseters compared to controls (Figure 6H, J, O; Table 2; p=0.034, T-test). Together, these results demonstrates that inactivation of *Mllt11* in NCCs and their neurogenic derivatives leads to disruptions in trigeminal nerve motor innervation of the masseter.

**Table 2.** Comparisons of trigeminal ganglion cellular composition at E12.5. SD, standard deviation. P-value derived from T-test.

| **Comparison** | **Control Mean ± SD; n** | **cKO Mean ± SD; n** | **P-value** |
| --- | --- | --- | --- |
| NF<sup>+</sup> fibres/mm<sup>2</sup> | 280.5 ± 65.8; 4 | 138.8 ± 114.8; 5 | 0.066 |
| BTX<sup>+</sup> points/mm<sup>2</sup> | 96.6 ± 20.88; 4 | 72.34 ± 20.10; 5 | 0.120 |
| NMJs/mm<sup>2</sup> | 79.11 ± 12.71; 4 | 38.89 ± 28.13; 5 | 0.034 |

## DISCUSSION

The aim of this study was to determine if the microtubule-interacting protein, Mllt11, plays a role in the formation of the trigeminal ganglion. Previous studies have demonstrated its requirement for neurogenic processes in different system through the mammalian nervous system including migration and neuritogenesis in lamination of the neocortex and the retina (Blommers et al 2023, Stanton-Turcotte et al 2022). Here we show that Mllt11 is required for the proper innervation of the masseter, suggesting a neuritogenic role in the peripheral nervous system. Using an NCC-specific ablation strategy we demonstrate that Mllt11 is necessary for maintaining an appropriate proportion of neural crest-derived and placode derived neurons in the trigeminal ganglia, and the innervation of one of its motor targets, the masseter.

In the trigeminal ganglion at E12.5 fate-mapped NCCs (tdTomato^+^ cells) represent a mix of both neurons and glia (Schwan cells), as both populations arise from the neural crest (Koontz et al 2023, Le Douarin & Smith 1988, Mendez-Maldonado et al 2020). We observed a reduction in NCC-derived Sox10^+^, but not placodal (Isl1/2^+^), upon *Wnt1*^*Cre2/+*^-mediated *Mllt11* ablation. However, *Mllt11* loss did not affect appear to overtly affect NCC migration, as evidence by largely normal patterns of NCC fate-mapped tdTomato^+^ cells populating trigeminal ganglia, suggesting it did not overtly affect NCC migration. Nor was the reduction of Sox10^+^ due to enhanced cell death. We therefore wondered if the deletion of *Mllt11* in NCC derivatives attenuated neuritogenesis, and thus investigated the innervation of one of the major targets of trigeminal nerve: the maseter. We found that *Mllt11* loss in peripheral ganglia resulted in reduced motor innervation of the masseter, as evident in the reduction of NF density and NF^+^/BTX^+^ (i.e. NMJs) in the muscle. NMJs are critical sites for communication between motor neurons and muscle fibers, and their proper assembly is essential for muscle contraction and motor control. Despite the reduction in innervation in the *Mllt11* mutants, we did not observe the any change in the density of BTX-rich regions, signifying they maintained post-synaptic assembly of the nicotinic acetylcholine receptor-dense apparatus. This is not surprising given that *Mllt11* expression is restricted to developing neurons (Yamada et al 2014) and our genetic strategy ablated it in peripheral neurons only. Kummer et al (Kummer et al 2004) demonstrated the cell-autonomous ability of skeletal muscle cells to form the post-synaptic motor assembly *in-vitro*, without a neuronal presence or signal. This is consistent with our *in vivo* observations of the maintenance of nitocotinic acetylcholine receptors in masseters of *Wnt1*^*Cre/+*^; *Mllt11*^*flox/flox*^ mutants despite the loss of invading neurites. The main role of the masseter is mastication, however, it plays a minor roles in facial expression, and speech (by stabilizing the mandible) in humans (Kent 2004). In mice, the role of the masseter is limited more to mastication and biting, providing the main muscle mass for these functions. Future research into the role of Mllt11 in peripheral nervous system development should examine the behavioural manifestations of the knockout in post-natal mice.

The reduced trigeminal motor innervation was principally attributed by a reduction in NCC-derived neurons and/or glia, due to the use of the *Wnt1*^*Cre2/+*^ driver mouse allele to delete *Mllt11*. In the most striking examples of masseter dysinnervation, we observed a near total absence of NF^+^ fibres. This suggests that placodal contributions to motor innervation from the trigeminal ganglion is limited and that a segregation of function is present between the crest and placodal populations in the ganglion, with the later playing a mainly sensory role. However, previous research suggests that cranial ganglia involved in strictly motor function (ciliary, oculomotor) with no sensory contributions are also composed of a combination of crest and placodal cells similar to the trigeminal (Lee et al 2003, Weglowski et al 2015), thus it is unlikely that a segregation of function is at play in the trigeminal ganglion. Future research should expand the analysis presented in this study to other cranial ganglia with differing precursor contributions, as well as examine the effect of *Mllt11* loss on sensory fibres.

With respect to possible molecular targets of Mllt11 action in nerve development, we previously identified α*/*β Tubulins, Actin and atypical Myosins as potential Mllt11-interaction targets in the fetal brain (Stanton-Turcotte et al 2022). The role of the cytoskeleton in NCC development is supported by work in chicken embryos showing that ß3Tub is expressed in pre-neuronal cells of the neural plate and dorsal neural tube, that is, the pre-migratory neural crest (Chacon & Rogers 2019). Interestingly, this expression coincides with the upregulation of NCC-specific markers such as *Sox10*. In contrast, *Mllt11* does not appear to be expressed in the pre-migratory or migratory NCCs. Instead, it is expressed later in differentiated neurons throughout the central and peripheral nervous systems, co-expressing with robust ß3Tub levels (Yamada et al., 2014). It is currently unclear whether Mllt11 promotes cytoskeletal dynamics in filopodia and growth cones to promote migration and axonogenesis, but its cytoskeletal association and neuritogenic phenotype in the cranial ganglia suggest that it may promote the formation of stabilized structures in the axon shaft (da Silva & Dotti 2002, Kapitein & Hoogenraad 2015). Similarly, in the central nervous system *Mllt11* is required for the formation of long rage axonal connections across the corpus callosum and the formation of complex dendritc arborizations in cortical pyramidal neurons (Stanton-Turcotte et al 2022). In addition, *Mllt11* promotes the migration of immature neurons in the cortex and retina, reflecting its expression in nascent neurons or neuroblasts (Blommers et al 2023, Stanton-Turcotte et al 2022). Given the restricted expression of Mllt11 in developing cranial ganglia and not crest cells reported herein, it is not surprising to find a mostly neuritogeneic role for *Mllt11* in the peripheral nervous system. Our findings are consistent with Mllt11 being a microtubule-interacting protein functioning to stabilize the cytoskeleton (Stanton-Turcotte et al 2022). As such the trigeminal ganglion neurons of cKO mice likely failed neuritogenesis, either due to disrupted neurite budding or polarization or elongation (da Silva & Dotti 2002). These processes rely on cytoskeletal reorganization and stabilization which are likely impaired in NCC-derived neurons lacking *Mllt11*, leading to reduction in the presence of distal motor fibres.

## CONCLUSIONS

This study demonstrates that the cytoskeletal-associated protein Mllt11 plays a role in the formation of the trigeminal ganglion. We show that *Mllt11* deficiency in NCC derivatives leads to alterations in the normal balance of cell types in the ganglion, and disrupts the motor innervation from the trigeminal ganglion. The most likely explanation of the phenotype being altered neuritogenesis of cranial nerves following *Mllt11* ablation. Cranial nerves, including the trigeminal nerve, play crucial roles in functions such as facial motor control and sensation. As such these findings have important implications for our understanding of cranial nerve development and motor function. Our study provides a foundation for future research on cranial ganglion development, and the investigation of mechansims regulating peripheral nerve innervation.

## ACKNOWLEDGEMENTS

We gratefully acknowledge funding from the Canadian Institutes of Health Research (CIHR PJT-388914), National Science and Engineering Research Council of Canada (RGPIN 03925-20), the Dalhousie University Faculty of Medicine Gladys Osman Estate, and the Killam Foundation for fellowship support to EW. We thank Sarah Whitehead for assistance with animal husbandry and Dr. Victor Rafuse for BTX reagents.

## REFERENCES

Blommers M, Stanton-Turcotte D, Iulianella A. 2023. Retinal neuroblast migration and ganglion cell layer organization require the cytoskeletal-interacting protein Mllt11. Dev Dyn 252: 305–19

Chacon J, Rogers CD. 2019. Early expression of Tubulin Beta-III in avian cranial neural crest cells. Gene Expr Patterns 34: 119067

Cordes SP. 2001. Molecular genetics of cranial nerve development in mouse. Nat Rev Neurosci 2: 611–23

D’Amico-Martel A, Noden DM. 1983. Contributions of placodal and neural crest cells to avian cranial peripheral ganglia. Am J Anat 166: 445–68

da Silva JS, Dotti CG. 2002. Breaking the neuronal sphere: regulation of the actin cytoskeleton in neuritogenesis. Nat Rev Neurosci 3: 694–704

Dinsmore CJ, Ke CY, Soriano P. 2022. The Wnt1-Cre2 transgene is active in the male germline. Genesis 60: e23468

Hamburger V. 1961. Experimental analysis of the dual origin of the trigeminal ganglion in the chick embryo. J Exp Zool 148: 91–123

Kaeser PF, Brodsky MC. 2013. Fourth cranial nerve palsy and Brown syndrome: two interrelated congenital cranial dysinnervation disorders? Curr Neurol Neurosci Rep 13: 352

Kapitein LC, Hoogenraad CC. 2015. Building the Neuronal Microtubule Cytoskeleton. Neuron 87: 492–506

Kent RD. 2004. The uniqueness of speech among motor systems. Clin Linguist Phon 18: 495–505

Koontz A, Urrutia HA, Bronner ME. 2023. Making a head: Neural crest and ectodermal placodes in cranial sensory development. Semin Cell Dev Biol 138: 15–27

Kummer TT, Misgeld T, Lichtman JW, Sanes JR. 2004. Nerve-independent formation of a topologically complex postsynaptic apparatus. J Cell Biol 164: 1077–87

Kurosaka H, Trainor PA, Leroux-Berger M, Iulianella A. 2015. Cranial nerve development requires co-ordinated Shh and canonical Wnt signaling. PLoS One 10: e0120821

Le Douarin NM, Smith J. 1988. Development of the peripheral nervous system from the neural crest. Annu Rev Cell Biol 4: 375–404

Lee VM, Sechrist JW, Luetolf S, Bronner-Fraser M. 2003. Both neural crest and placode contribute to the ciliary ganglion and oculomotor nerve. Dev Biol 263: 176–90

Mendez-Maldonado K, Vega-Lopez GA, Aybar MJ, Velasco I. 2020. Neurogenesis From Neural Crest Cells: Molecular Mechanisms in the Formation of Cranial Nerves and Ganglia. Front Cell Dev Biol 8: 635

Porter AG, Janicke RU. 1999. Emerging roles of caspase-3 in apoptosis. Cell Death Differ 6: 99–104

Stanton-Turcotte D, Hsu K, Moore SA, Yamada M, Fawcett JP, Iulianella A. 2022. Mllt11 Regulates Migration and Neurite Outgrowth of Cortical Projection Neurons during Development. J Neurosci 42: 3931–48

Sudiwala S, Knox SM. 2019. The emerging role of cranial nerves in shaping craniofacial development. Genesis 57: e23282

Traboulsi EI. 2004. Congenital abnormalities of cranial nerve development: overview, molecular mechanisms, and further evidence of heterogeneity and complexity of syndromes with congenital limitation of eye movements. Trans Am Ophthalmol Soc 102: 373–89

Trainor PA. 2014. Neural crest cells : evolution, development, and disease. Amsterdam; Boston: Elsevier/AP, Academic Press is an imprint of Elsevier. xviii, 469 pages pp.

Weglowski M, Wozniak W, Piotrowski A, Bruska M, Weglowska J, et al. 2015. Early development of the facial nerve in human embryos at stages 13-15. Folia Morphol (Warsz) 74: 252–7

Yamada M, Clark J, Iulianella A. 2014. MLLT11/AF1q is differentially expressed in maturing neurons during development. Gene Expr Patterns 15: 80–7

